# Atypical gaze patterns in autism are heterogeneous across subjects but reliable within individuals

**DOI:** 10.1101/2021.07.01.450793

**Authors:** Umit Keles, Dorit Kliemann, Lisa Byrge, Heini Saarimäki, Lynn K. Paul, Daniel P. Kennedy, Ralph Adolphs

## Abstract

People with autism spectrum disorder (ASD) have atypical gaze onto both static visual images ^1,2^ and dynamic videos ^3,4^ that could be leveraged for diagnostic purposes ^5,6^. Eye tracking is important for characterizing ASD across the lifespan ^7^ and nowadays feasible at home (e.g., from smartphones ^8^). Yet gaze-based classification has been difficult to achieve, due to sources of variance both across and within subjects. Here we test three competing hypotheses: (a) that ASD could be successfully classified from the fact that gaze patterns are less reliable or noisier than in controls, (b) that gaze patterns are atypical and heterogeneous across ASD subjects but reliable over time within a subject, or (c) that gaze patterns are individually reliable and also homogenous among individuals with ASD. Leveraging dense eye tracking data from two different full-length television sitcom episodes in a total of over 150 subjects (N = 53 ASD, 107 controls) collected at two different sites, we demonstrate support for the second of these hypotheses. The findings pave the way for the investigation of autism subtypes, and for elucidating the specific visual features that best discriminate gaze patterns — directions that will also inform neuroimaging and genetic studies of this complex disorder.

## Results

### Participants and Stimuli

Autism spectrum disorder (ASD) is widely recognized to be a complex and heterogeneous disorder that has so far defied any simple diagnostic markers or cognitive mechanisms ^9^. Yet there is universal agreement that processing of social stimuli is prominently atypical (a core component of the diagnosis) ^10,11^ and studies suggest that many different features contribute. Perhaps the most robust general finding is that people with ASD do not fixate faces typically, including expressive features (eyes, mouth) within faces, although this depends on context ^1,3,12^. Another robust finding is that infants with ASD do not gaze typically at biological motion stimuli ^13^. Other theories propose atypical attention to visual stimuli that involve other people’s direction of eye gaze ^14^, prediction ^15^, geometric patterns ^16^, reward value ^17^, or imitation engagement ^18^. Our own work has similarly argued that atypical gaze in ASD is distributed across a broad range of visual features ^2^. This background motivated our choice of stimuli – a rich and highly social television sitcom (“The Office”; NBC Universal) that we have found successful in our prior neuroimaging work on ASD 19,20.

To discover biological markers that might distinguish people with ASD, we analyzed eye tracking data from a sample of high-functioning adults with ASD and typically developed controls (TD) matched on age-, sex- and Full Scale IQ (Table 1). To best distinguish our three hypotheses, we aimed to estimate each subject’s reliability in gaze most precisely. For this reason, we incorporated the following features into our study. First, we selected high-functioning adults who could follow instructions and could enjoy watching the stimuli, reducing contributions of general inattention. Second, we tested subjects at two different sites (Caltech, and Indiana University, IU) to ensure that findings generalize across labs and subject samples. Third, we used two different video episodes to test within-subject reliability across stimuli.

**Table 1.**
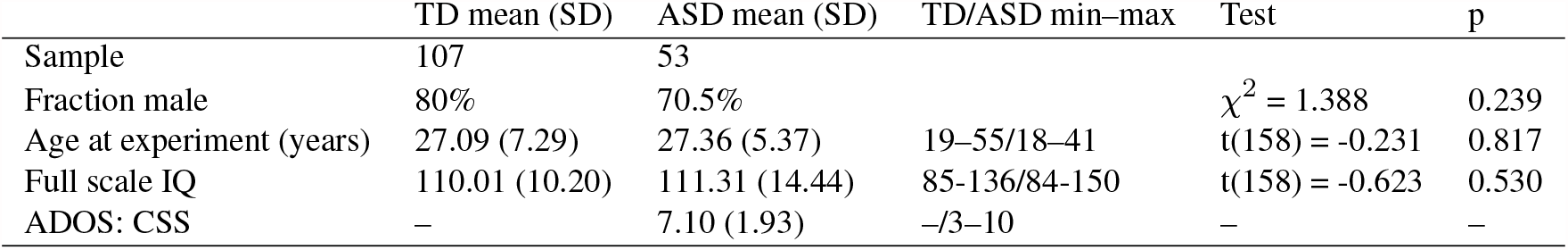
Participants’ demographic information, IQ, and ADOS scores (for ASD participants)

|  | TD mean (SD) | ASD mean (SD) | TD/ASD min–max | Test | p |
| --- | --- | --- | --- | --- | --- |
| Sample | 107 | 53 |  |  |  |
| Fraction male | 80% | 70.5% | | $\chi^2 = 1.388$ | 0.239 |
| Age at experiment (years) | 27.09 (7.29) | 27.36 (5.37) | 19–55/18–41 | $t(158) = -0.231$ | 0.817 |
| Full scale IQ | 110.01 (10.20) | 111.31 (14.44) | 85–136/84–150 | $t(158) = -0.623$ | 0.530 |
| ADOS: CSS | – | 7.10 (1.93) | –/3–10 | – | – |

### Eyetracking to social features

We first quantified gaze differences between ASD and controls by calculating the percentage of time spent looking at several predefined visual features in the videos, such as faces, eyes, or mouth (“Areas of Interest”, AOI; see Methods). As expected, the ASD group looked less at faces, but more at other body parts or non-social content (inanimate objects and/or background), and also looked less at eyes within faces (Cohen’s d = 0.807 in Episode A and d = 0.750 in Episode B for faces, *p* < 0.001, bootstrap test; Figs. 1A, 1B, and 1C). This feature-based analysis was complemented by an alternative analysis using gaze heatmaps. A reference gaze heatmap was created for each 1-second time bin of videos by combining data from all TD controls. An individual’s gaze similarity to this reference visual salience was quantified as the Pearson correlation between individual heatmaps and the reference heatmap in 1-second time bins (a leave-one-subject-out approach was used to prevent bias for TD individuals, see Methods). We found that the heatmap correlation was significantly higher in the TD group than in the ASD group (d = 0.859 in Episode A, d = 0.752 in Episode B, *p* < 0.001, bootstrap test; Fig. 1D). A complementary analysis based on constructing a reference heatmap from all ASD participants in a leave-one-out fashion corroborated these results (d = 0.840 in Episode A, d = 0.729 in Episode B, *p* < 0.001, bootstrap test; Fig. 1D): the TD group was still more strongly correlated to this new reference heatmap than the ASD group, already providing some initial evidence against hypothesis 3 (we provide stronger evidence below). The reference heatmap based on ASD individuals was also highly correlated with the reference heatmap based on TD controls (0.977 ± 0.016 in Episode A; 0.969 ± 0.021 in Episode B, mean ± standard deviation across 1-second time bins within each episode). A more detailed analysis based on resampling of different duration epochs from videos and bootstrap resampling of individual participants was used to examine the effect of reducing data size on assessed group differences. We found that the significant group differences can be observed even with 1-minute samples of the videos (Supplementary Table 1).

**Figure 1.**
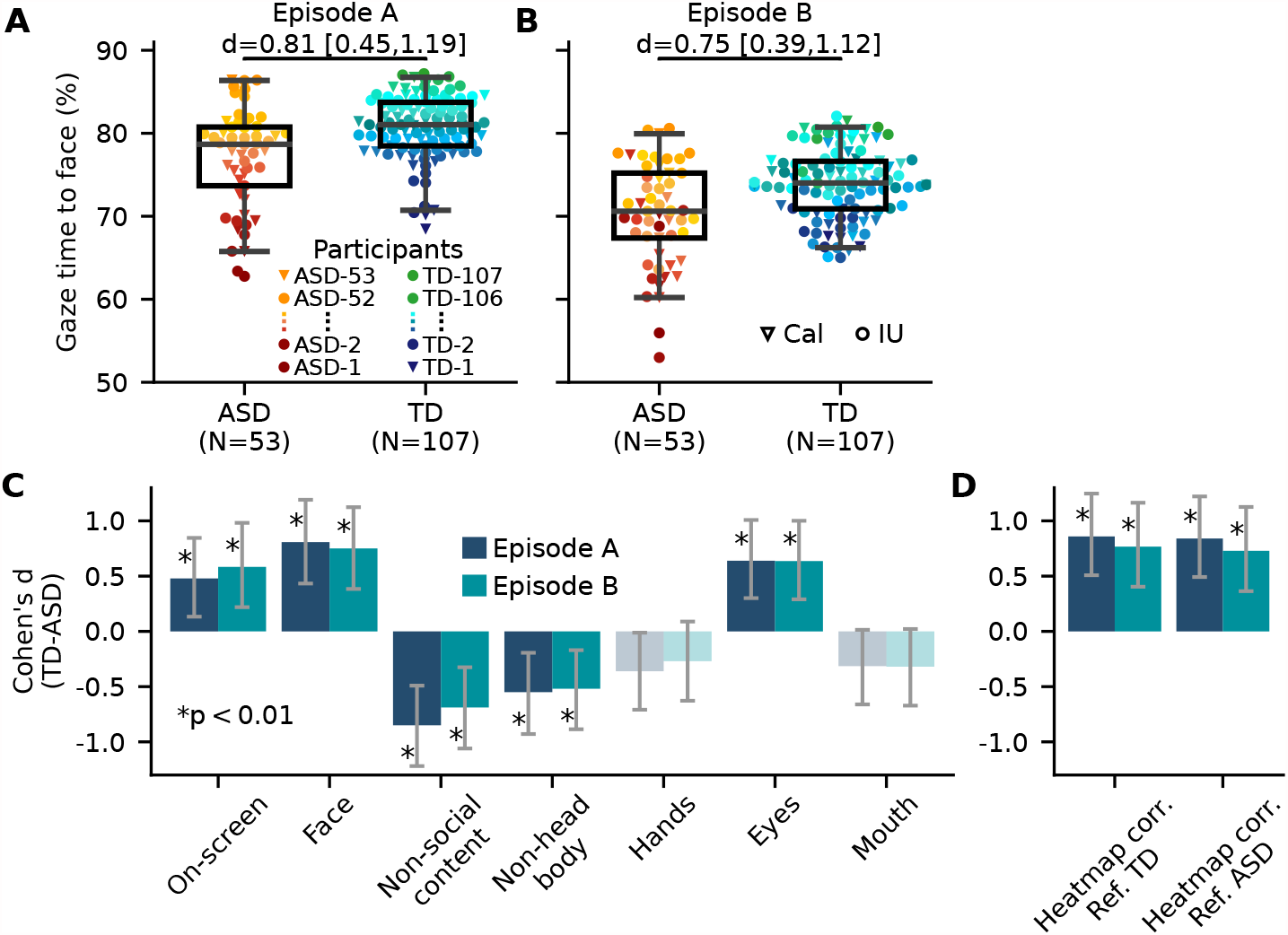
Eye tracking demonstrates reliable gaze differences to features of videos. (A and B) ASD versus TD comparison in their percentage of total gaze time to faces in two separate videos. Error bars span the 2.5^*th*^ to 97.5^*th*^ percentiles, boxes span the 25^*th*^ to 75^*th*^ percentiles, and horizontal black lines indicate medians. Effect size of the difference between groups (Cohen’s d) is given on top of the figures. Individuals were marked with different red-yellow spectrum colors based on their percentage of gaze time to faces in Episode A and the same subject-wise colors were used for Episode B (and also for Fig. 2). Inverted triangles: subjects from Caltech site; circles: subjects from IU site.(C) Effect size of the differences between groups in their percentage of total gaze time to several regions of interest in two separate videos. The bar height indicates Cohen’s d and error bars indicate its bootstrap confidence interval. Saturated colors, asterisks, and p-values indicate the statistical significance of Cohen’s d (*p* < 0.05, assessed with bootstrap tests, and corrected for multiple comparisons via FDR); desaturated color indicates nonsignificant differences.(D) Effect size of the differences between groups in their average correlation with reference gaze heatmaps created by either combining all TD controls (Ref. TD) or all ASD individuals (Ref. ASD). Same format as (C).

### Within-subject reliability

We next examined whether the discriminating differences between the groups (Fig. 1C, 1D) were driven by noisier (less reliable) gaze patterns in ASD. We reasoned that if people with ASD have less reliable gaze patterns, then the rank-order among ASD individuals in their gaze time to various AOIs and among heatmap correlations should reflect this variability when examining within-subject data between two episodes or across epochs of videos. An initial test comparing rank-order correlations between two different video episodes showed that this prediction was incorrect: those subjects who look least at specific features (e.g., faces) in one video, also do so reliably in a second video (Fig. 2). A more detailed analysis based on resampling of 1-minute epochs from each video’s data confirmed equivalent and high within-subject reliability in the ASD and TD groups (all correlations significantly different from zero; Spearman’s *ρ* > 0.410, *p* < 0.05, bootstrap test, FDR corrected, Figs. 2C, 2F, and 2I). Furthermore, correlation values did not differ between subject groups (*p* = 0.842 for faces; *p* = 0.731 for eyes; *p* = 0.678 for heatmap correlation; bootstrap test, FDR corrected). In an additional analysis, we examined the correlation values using different duration resampling epochs in a range of 10 minutes to 10 seconds. We found that although correlation values decreased with decreasing epoch duration, as would be expected, there was no significant difference between the values assessed for ASD and TD groups (*p* > 0.310, bootstrap test; Supplementary Table 2). Furthermore, these findings were robust with respect to quality control: when more stringent subject exclusions were applied based on the eye tracking calibration that preceded each session, this decreased our sample size but did not substantively changed the pattern of results (Supplementary Fig. 1). These results argue that the gaze patterns of individuals with ASD are as reliable as those of TD controls, evidence against hypothesis 1.

**Figure 2.**
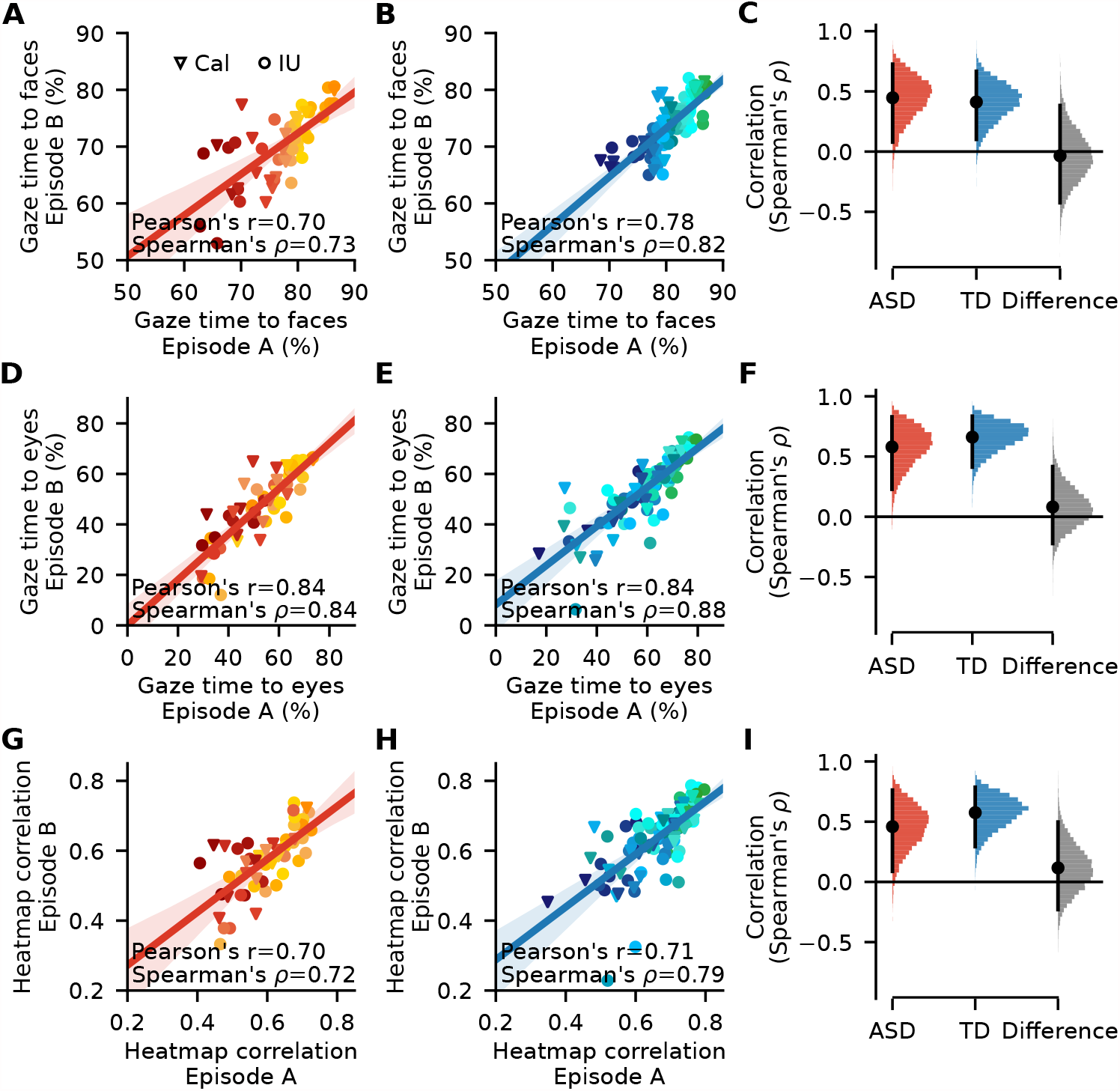
High within-subject reliability in ASD. (A, B) Individual participants’ percentage of gaze time to faces is plotted from two separate videos. Line: Pearson correlation and bootstrapped CI. Individual subjects are denoted by red-yellow (ASD) or dark-light blue (TD) spectrum colors encoding the percentage of gaze time to faces in Episode A (panels A, B) and the same subject-wise color codes were used for other panels (panels D,E, G, and H). Triangular (circular) markers indicate subjects from the Caltech (IU) site. (D, E) Individual participants’ percentage of gaze time to eyes is plotted from two separate videos. Same format as (A, B). (G, H) Individual participants’ average gaze heatmap correlation with TD-reference gaze heatmaps. Same format as (A, B). (C,E,I) Resampling analysis based on 1-min epoch from the videos and bootstrap resampling of individual participants. Color scheme is the same as in Fig. 1.

To corroborate these findings, we asked whether individuals can be distinguished from other participants based on their patterns of gaze alone. For this analysis, we used a “gaze fingerprinting” procedure ^21,22^ which provided a multivariate analysis approach complementing the univariate approach above. We examined the identification of individuals based on the similarity between their gaze patterns across two episodes, quantified using the distribution over eight gaze features: the percentage of time spent looking at screen, faces, non-social content, non-head body parts, hands, eyes, mouth, and heatmap correlations with TD-reference heatmaps. This provided a fingerprinting identification accuracy of 32.1% (17/53) and 29.0% (31/107) for ASD and TD, respectively, both significantly above chance (*p* < 0.001, permutation test; chance was 5.7% (3/53) and 2.8% (3/107) from the 95^*th*^ percentile of an empirical null distribution). A more robust analysis based on resampling of 10-minute epochs and bootstrap sampling of individual participants from video data provided similar accuracies (33.3% for ASD and 34.0% for TD group; *p* < 0.001, permutation test, FDR corrected). Furthermore, groups did not differ in identification accuracy (mean[ASD-TD] = 0.7%, *p* = 0.99, bootstrap test).

To test how fingerprinting identification accuracy scales with duration of the video we gradually reduced the sampling epoch and found that the identification accuracy for either group was significantly higher than chance already for 1-minute epochs (14.7%, *p* = 0.021 for ASD and 13.3%, *p* = 0.010 for TD group), but dropped to chance level for 30-second epochs (11.0%, *p* = 0.071 for ASD and 10.0%, *p* = 0.043 for TD group, see Supplementary Table 3). We also examined whether there might be a subset of the eight gaze features that might carry sufficient information for fingerprinting. We removed non-social content, non-head body parts, and hand-looking times from the feature set because they are highly correlated with face-looking time. We also removed mouth-looking time because of its high-correlation with eye-looking time. Gaze fingerprinting analysis based on the four remaining features (percentage of on-screen, face- and eye-looking time, heatmap correlations) yields nearly the same identification accuracies as the full set of features (31.1% for ASD and 33.0% for TD group for 10-minute epochs, *p* < 0.001; 14.8% for ASD and 14.5% for TD group for 1-minute epochs). None of these additional analyses revealed any significant difference between the identification accuracy estimated for ASD and TD groups (*p* > 0.90, bootstrap test), consistent with our initial rank-order correlation analyses. Taken together, these findings demonstrate substantial within-subject reliability in gaze that is equivalent in ASD and TD, and that is distributed across multiple visual features.

### Within-group heterogeneity

Figure 2 confirms substantial within-subject reliability despite considerable between-subject variability: some individuals always gaze at faces, while some reliably rarely do. Supporting our hypothesis 2, this pattern of results suggests that dichotomous classification of individuals as ASD versus TD should be difficult, but that a data-driven clustering might still reveal subgroups within the ASD group. To examine the supervised dichotomous classification of ASD vs. TD, we used a Gaussian Naive Bayes classifier. To robustly estimate classification accuracy, we re-sampled a 10-minute epoch from video data, randomly held-out 15 ASD and 15 TD individuals, trained a classifier in the remaining participants’ data and finally assessed the classification accuracy based on data from held-out participants. As the fingerprinting procedure above indicated, individuals can be reliably identified from their gaze features based on 10-minute gaze data. This resampling and model estimation process was repeated 5,000 times. We found an overall above chance classification accuracy (0.80 ± 0.077; mean ± SD across CV iterations; see classification confusion matrix, Fig. 3A). When we examined the frequency of correct classification and misclassification of individual participants across 5,000 CV iterations of the classifier, we found that 81 participants, 19 with ASD, were correctly classified more than 90% of the time across cross-validated iterations. However, 23 participants, 11 with ASD, were misclassified more than 90% of the time (see Fig. 3B). This confirms, as expected, that the heterogeneous gaze patterns of ASD individuals preclude diagnostically robust classification. Control analyses using different time bins and different gaze parameters did not change these results (see Methods). Moreover, classification accuracy varied considerably across time windows of the video, as expected given the large variability of features and gaze onto them.

**Figure 3.**
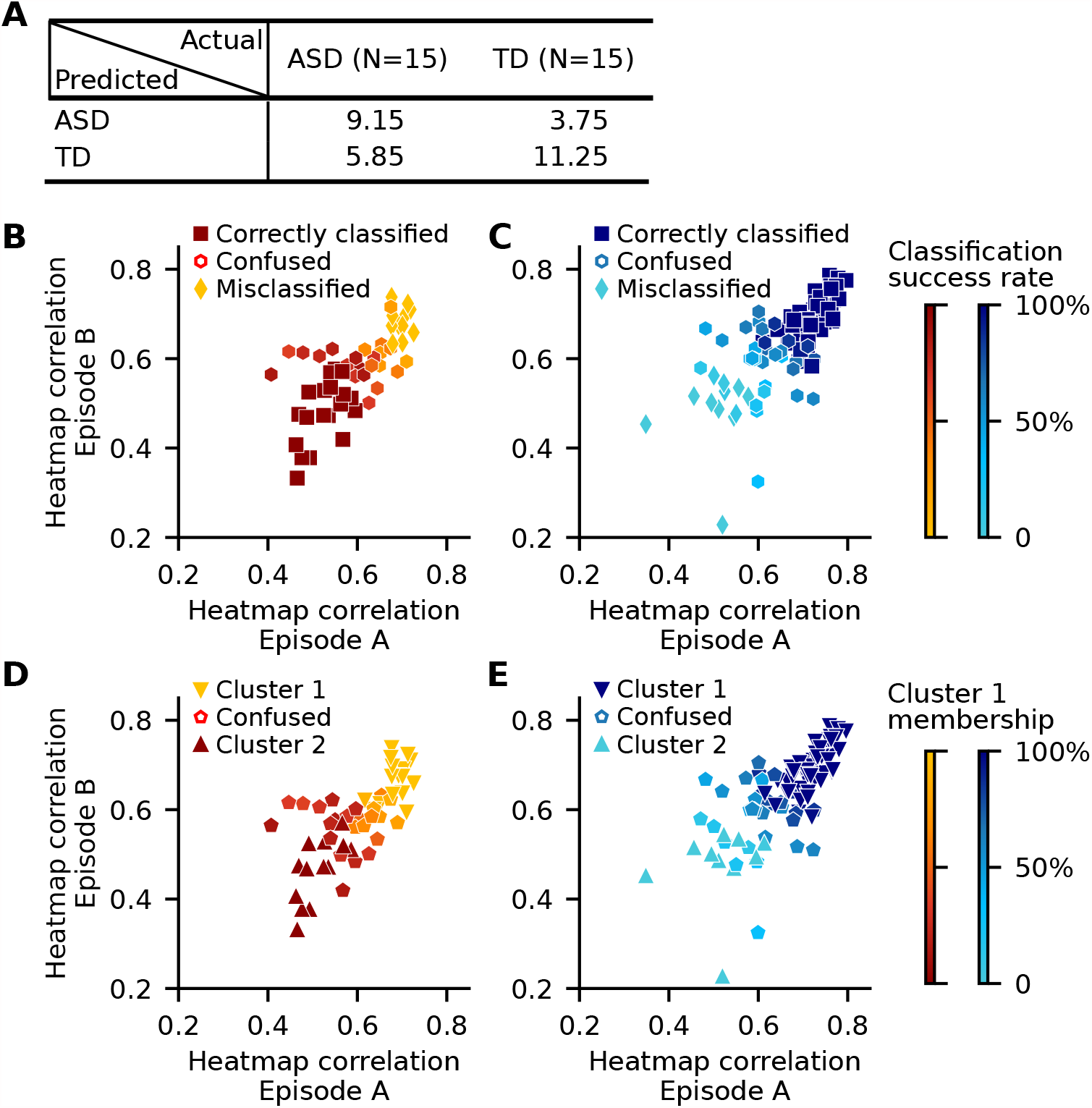
Classification and clustering of participants. (A, B) Classification confusion matrix for Gaussian naive Bayes classifier.(C, D) Correctly classified and mis-classified participants across 5,000 crossvalidation iterations of a Gaussian naive Bayes classifier. Participants denoted by square (diamond) markers were correctly classified (misclassified) more than 90% of the time across CV iterations. Participants shown with hexagon markers were classified with a frequency lower than 90% (i.e., confused as ASD or TD in different iterations). (B) ASD participants. (C) TD participants. Color bar depicts at what frequency an individual participant was classified correctly. (D,E) Unsupervised clustering of participants into subgroups using a Gaussian mixture model procedure. The clustering procedure identified two groups of participants (indicated as Cluster 1 and 2) based on each participant’s gaze similarity to the reference TD heat map. (D) ASD participants. (E) TD participants. Color bar depicts at what frequency an individual participant was partitioned to Cluster 1 (i.e., Cluster 1 membership). Although the analyses were based on the time course of heatmap correlations within 1-second time bins and resampling of 10-minute epochs, here we used the same format as in Figs. 2G and 2H to visualize the classification and clustering results.

### Unsupervised clustering

Could the individual-level reliability in gaze patterns be used to discover subtypes of ASD? We next used a Gaussian mixture model to automatically assign participants into subgroups based on each participant’s gaze patterns. Using a moving window of 10-minute epochs across video episodes, the number of subgroups were learned from data for 33 separate times. Notably, in all these epochs, the model identified two subgroups; one larger group (Cluster 1; n = 109.21 ± 5.96, mean ± SD across 33 epochs; 26.06 ± 2.89 ASD with 61 TD and 12 ASD common across all 33 epochs) and one smaller group (Cluster 2; n = 50.79 ± 5.96 with 26.94 ± 2.89 ASD; 8 TD and 10 ASD common across all 33 epochs; Figs. 3D, 3E). These numbers suggest that Cluster 1 across epochs captured largely TD controls whereas Cluster 2 captured those ASD and TD individuals who deviated from the majority of TD controls. Overall, the clustering results corroborated the prior classification result that ASD individuals cannot be regarded as a homogenous group, but rather consisted of at least two separate subgroups of ASD individuals, some of them reliably similar to TD controls, and others reliably dissimilar.

In conclusion, we find support for our hypothesis 2: gaze patterns of people with ASD are strongly heterogeneous yet reliable within subjects. Our clustering analysis points to two possible subgroups: those with gaze similar to TD controls, and a heterogeneous subgroup with reliably atypical gaze. This pattern is reminiscent of neuroimaging results we ^19^ and others ^23^ have documented previously, and leaves open the discovery of homogenous ASD subtypes if larger sample sizes can be analyzed in future studies. The systematic differences in how people with ASD fixate visual stimuli compared to TD controls would be expected to translate to differences also in evoked BOLD-fMRI activations ^24^, and may be an endophenotype reflecting substantial genetic effects ^12,22^. While genetics, gaze, and neuroimaging data will all need to be put together for a comprehensive mechanistic explanation of ASD, eye tracking will continue to have strong practical advantages, given current technology to collect it at home from laptop-based ^25^ or smartphone-based cameras ^8^. Such in-home collection could be used in order to achieve the larger sample sizes required to further test both for longterm longitudinal stability within individuals, as well as to explore possible subtypes among them.

## Methods

### Participants

Data used in this study were collected as part of a larger study that included extensive behavioral assessment, neuroimaging, and eye tracking. Here we present only the eye tracking data (collected outside the scanner), together with a subset of the behavioral assessment data. We initially recruited a total of 188 individuals, 72 of whom were high-functioning adults with a DSM-5 diagnosis of ASD (mean age 27.1; range 18–46; 20 female; 22 tested at the Caltech site) and 116 of whom were typically developed, community-dwelling controls (TD controls; mean age 27.2; range 19–55; 33 female; 34 tested at the Caltech site). 185 participants watched Episode A (see details of stimuli below; 68 ASD, 117 TD), and 178 participants watched Episode B (68 ASD, 110 NT). We limited our analyses to those 174 participants who watched both Episode A and B (64 ASD, 110 NT). We subsequently excluded any individual with excessive missing data, defined as less than 50% of data in either one or both episodes (10 ASD, 2 TD). Furthermore, one ASD and one TD participant exhibiting mean heatmap correlation values exceeding 4 SD from the mean of all participants were excluded to produce a more normal distribution (but results remain essentially unchanged with their inclusion). The final sample consisted of 53 participants with ASD (17 Caltech site) and 107 TD participants (33 Caltech site) of similar age, sex ratio, and Full-Scale IQ (Table 1). All conclusions held across data from each of the two sites separately; we therefore pooled subjects in all analyses we present here.

### Stimuli

Participants freely watched two episodes of the television sitcom “The Office” (NBC Universal, originally aired in 2005). Episode 1 of Season 1 (22 min, called Episode A here) was viewed in three separate parts, one shortly after the next one (part 1, 6 min 58 sec; part 2, 8 min 30 sec; part 3, 6 min 28 sec). Episode 4 (21 min, called Episode B here) was paused at times and participants were briefly asked to verbally respond to social comprehension questions, as described in Byrge, Dubois, et al. ^19^, but these moments were excluded from the analysis. Both episodes were viewed on the same day separated by a break.

### Eye tracking

Participants were comfortably seated in front of a Tobii TX300 eye tracker at a distance of approximately 65 cm from the screen (a movable 23”, 1920 *×* 1080 widescreen monitor). The eye tracker provided calibrated gaze data at 300Hz (0.4 degrees spatial resolution). We focused our analyses on the raw gaze data; results are similar when fixations are derived as the data input. Prior to starting the videos, participants carried out a 9-point calibration on the screen, followed by a 9-point validation in which the gaze error to the 9 calibrated locations was quantified; the 9 target dots spanned the full extent of the screen. For Episode A, the calibration-validation procedure was repeated prior to viewing each of three parts. For Episode B, the procedure was completed once at the very beginning of the episode. Quantitative accuracy results were immediately displayed to the experimenter, who would adjust the screen and/or participant and redo the calibration-validation procedure if any of the points had *>* 1.5 degrees of error. A computer error resulted in loss of some of the validation data.

### Automatic segmentation of frames to area of interests (AOIs)

Human body parts in each frame of videos were detected by using a pre-trained neural network model provided in the DensePose module in the Detectron2 deep learning software ^26,27^. The body parts provided by the DensePose model were merged to obtain three main parts of the body for our interest: the head areas, hands, and non-head body area. The regions within a frame where no body parts were detected were taken as non-social content. Face areas and five facial keypoints, including two eyes, nose, and two sides of the mouth, within each frame were detected by using the RetinaFace face detector model provided in the InsightFace deep face analysis toolbox ^28^.

### Estimation of effect sizes in group differences

Cohen’s d was used to measure the effect size of difference between ASD and TD groups. A bootstrap procedure was used to robustly estimate Cohen’s d and its confidence interval. Individual participant’s gaze time percentages or heatmap correlation values were randomly sampled for 10,000 times with replacement separately for each group, and Cohen’s d between groups was computed at each iteration. The mean Cohen’s d for each comparison was obtained by averaging the d values across bootstrap iterations. Confidence interval of Cohen’s d was taken as 2.5^*th*^ and 97.5^*th*^ percentiles of the bootstrap distribution.

### Quantifying spatiotemporal gaze patterns using heatmaps

Temporally-binned heatmaps were used to assess each subject’s gaze similarity with a reference group’s gaze (leaving one subject out for each group gaze calculation, if that subject was a member of the respective reference group). A gaze heatmap was built for each subject for each time bin by convolving each gaze point in the time bin with a two-dimensional isotropic Gaussian with a standard deviation of 0.5 degrees of visual angle. One degree of visual angle corresponded to 41 pixels on the screen in our study. A reference gaze heatmap was built for each time bin by applying the same Gaussian convolution on aggregate gaze data from all subjects in a group (TD for the TD-ref heat map; ASD for the ASD-ref heat map; cf. Fig. 1D). The reference gaze heatmaps thus reflected the visual salience of the videos for that subject group (both TD-ref and ASD-ref heat maps were highly correlated; see main text). To estimate how an individual’s gaze aligned with this reference visual salience at each time bin, we calculated the correlation between the individual’s gaze heatmap and the reference heatmap for each time bin. To calculate correlation coefficients between heatmaps, each heatmap was first converted to a single vector of values, and then the Pearson correlation between these two vectors was calculated.

### Gaze fingerprinting

To estimate whether an individual’s gaze to visual features was reliable across different epochs of videos, we used a gaze fingerprinting approach. In this approach, each individual’s gaze data within a time bin were represented as an eight-dimensional gaze vector (the percentage of time spent looking at screen, faces, non-social content, non-head body parts, hands, eyes, mouth, and heatmap correlations with TD-reference heatmaps). In the identification procedure, given a gaze vector of a subject in a time bin, we calculated the pairwise L1-distances between this gaze vector and all gaze vectors from all subjects (including themselves) from another time bin. The subject identity was predicted based on the minimum pairwise distance. This procedure was repeated for each individual and the successful identifications were counted. Prior to computing distances, each feature channel in the gaze vectors was standardized to zero mean and unit variance across subjects within each time bin to control for range and variance differences between features. The statistical significance of identification accuracy was computed by a permutation test with 10,000 iterations. At each iteration of the permutation, identities were shuffled across gaze vectors, and then the fraction of successful identifications computed. The obtained empirical null distribution of identification accuracy was used to assess the p-value of the empirical accuracy.

### Gaussian mixture model

For an unsupervised partitioning of subjects into subgroups, we trained a variational Bayesian Gaussian mixture model ^29^. This approach allowed us to learn the number of subgroups (i.e., clusters) from data automatically. To obtain multiple estimates of the number of clusters over different epochs of videos, we first combined first gaze data from the two video episodes into a single 43 minute dataset. Using a moving window of 10-minute length with step size of 1-minute, the gaze data were resampled 33 times. Thus, at each epoch, each subject was represented by a 600-dimensional vector, where each dimension was the correlation between an individual’s gaze heatmap and the TD-reference heatmap within a 1-second block. These subject-wise 600-dimensional vectors were concatenated across subjects to obtain a two-dimensional matrix with dimensions of the number of subjects by 600. A Gaussian mixture model was fit to this matrix to reveal underlying clusters based on gaze similarities between subjects. The model was initiated to identify five clusters, but returned two large clusters (each containing more than 30 subjects) and three small clusters (each containing 1 or 2 subjects only). Individuals in these three small clusters were then assigned to one of the large clusters based on the minimum Euclidean distance between an individual’s 600-dimensional gaze vector and the mean gaze vector representing the cluster center. To perform this analysis, we used the implementation of variational Bayesian Gaussian mixture models in the Python machine learning library scikit-learn ^30^.

## Acknowledgements

This research was supported in part by grant R01MH110630 from NIMH (DPK/RA), the Simons Foundation Autism Research Initiative (RA), the Simons Foundation Collaboration on the Global Brain (542951; UK), and a Della Martin Fellowship (DK). We are grateful for Tim Armstrong and Steven Lograsso for help with data collection, and to all of our participants and their families for their participation in this time-intensive study. We thank David Kahn for useful discussions regarding the project.

## Author contributions

DPK, LB, and RA designed the study. LKP and DPK recruited and assessed subjects. DK, SL, and Indiana University personnel collected data. UK pre-processed data, ensured data quality, and conducted all analyses. RA and UK drafted the paper and all co-authors provided extensive discussions and edits.

## Competing interests

The authors declare no competing interests.

## Supplementary Information

**Supplementary Figure 1.**
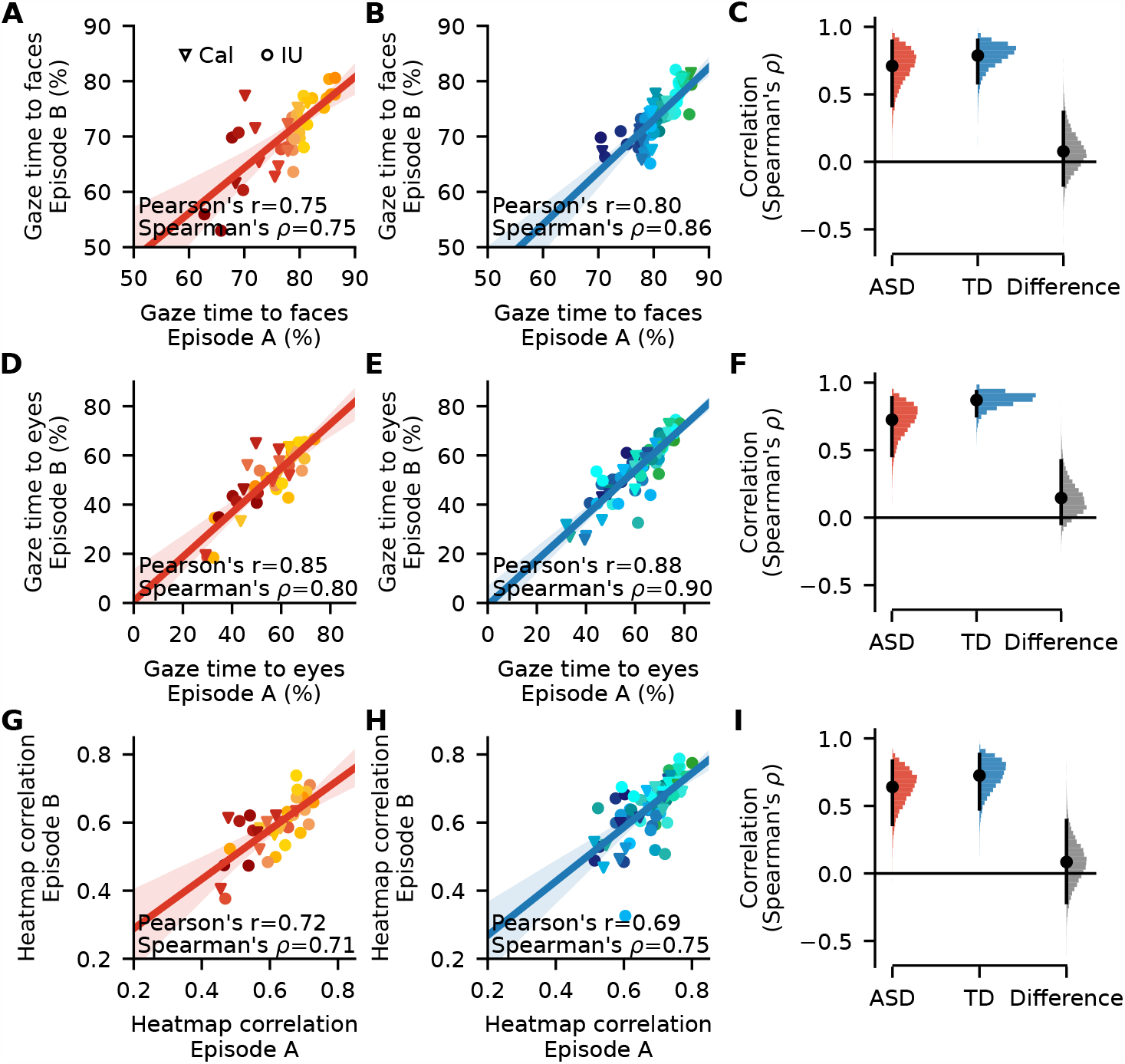
Gaze reliability within subjects for more stringently selected subjects. All panels show the same analyses as in Fig. 2, except for a more stringently selected subset of subjects (ASD N = 38; TD N = 76). 10-minute time bins were used to compute the values in the rightmost column (rather than 1-minute in Fig. 2). For this more stringent analysis, participants were excluded if their 9-point calibration distance was *>* 3SD from the group mean calibration distance. Calibration distances were calculated as the mean of each subject’s deviation from each of the 9-calibration points. Note that many participants were excluded due to missing calibration data rather than known poor calibration quality. Format is the same as in Fig. 2.

**Table 1.** Effect size of the differences (quantified with Cohen’s d) between groups in their percentage of total gaze time on-screen, faces, eyes, and in their average correlation with TD-reference gaze heatmaps. Cohen’s d between the groups (TD-ASD) were computed within randomly sampled epochs of videos (duration given in rows) and then averaged across 10,000 resampling iterations. Values in parentheses show significance level (bootstrap test, FDR corrected for multiple comparisons within each epoch duration). This table complements Figs. 1C and 1D using a resampling procedure that examines the effect of reducing data size on estimated group differences. Asterisk denotes *p <* 0.001.

| Epoch duration | Gaze features |  |  |  |
| --- | --- | --- | --- | --- |
|  | On-screen | Faces | Eyes | Heatmap Corr. |
| 10 mins | 0.470 (0.010) | 0.751 (*) | 0.618 (*) | 0.777 (*) |
| 5 mins | 0.452 (0.020) | 0.711 (0.002) | 0.611 (0.002) | 0.750 (*) |
| 2 mins | 0.411 (0.044) | 0.634 (0.003) | 0.579 (0.003) | 0.701 (*) |
| 1 min | 0.375 (0.082) | 0.550 (0.018) | 0.554 (0.018) | 0.658 (0.003) |
| 30 secs | 0.343 (0.131) | 0.480 (0.060) | 0.515 (0.041) | 0.609 (0.030) |
| 20 secs | 0.313 (0.178) | 0.434 (0.099) | 0.489 (0.054) | 0.569 (0.044) |
| 10 secs | 0.288 (0.242) | 0.369 (0.175) | 0.446 (0.104) | 0.509 (0.104) |

**Table 2.** The Spearman correlation among individuals within a group in their gaze time to various AOIs and in their gaze heatmap correlation with TD-reference heatmap. Correlation values were computed between two randomly sampled epochs of videos (duration given in columns) and then averaged across 10,000 resampling iterations. Values in parentheses show significance level (bootstrap test, FDR corrected for multiple comparisons within each epoch duration). Correlation values did not differ between ASD and TD groups in any of the considered gaze feature or epoch duration (*p >* 0.310). This table complements analyses provided in Figs. 2E, 2F, and 2I for different duration resampling epochs. Asterisk denotes *p <* 0.001.

| Gaze feature | Group | Resampling epoch duration |  |  |  |  |  |  |
| --- | --- | --- | --- | --- | --- | --- | --- | --- |
|  |  | 10 mins | 5 mins | 2 mins | 1 min | 30 secs | 20 secs | 10 secs |
| On-screen | ASD | 0.643(*) | 0.627(*) | 0.552(0.007) | 0.494(0.028) | 0.435(0.059) | 0.405(0.106) | 0.350(0.230) |
|  | TD | 0.790(*) | 0.769(*) | 0.694(*) | 0.629(0.001) | 0.559(0.005) | 0.522(0.019) | 0.459(0.079) |
|  | TD-ASD | 0.148(0.310) | 0.142(0.402) | 0.142(0.557) | 0.135(0.656) | 0.125(0.743) | 0.117(0.813) | 0.109(0.860) |
| Faces | ASD | 0.697(*) | 0.663(*) | 0.554(0.007) | 0.450(0.037) | 0.335(0.148) | 0.281(0.279) | 0.197(0.530) |
|  | TD | 0.767(*) | 0.684(*) | 0.537(*) | 0.410(0.028) | 0.297(0.148) | 0.239(0.279) | 0.163(0.530) |
|  | TD-ASD | 0.069(0.565) | 0.021(0.929) | -0.017(0.903) | -0.040(0.842) | -0.038(0.867) | -0.042(0.863) | -0.034(0.888) |
| Eyes | ASD | 0.775(*) | 0.733(*) | 0.663(*) | 0.585(0.007) | 0.492(0.059) | 0.440(0.106) | 0.351(0.268) |
|  | TD | 0.863(*) | 0.824(*) | 0.746(*) | 0.661(0.001) | 0.561(0.005) | 0.499(0.043) | 0.395(0.230) |
|  | TD-ASD | 0.088(0.388) | 0.091(0.460) | 0.084(0.612) | 0.077(0.731) | 0.069(0.812) | 0.060(0.861) | 0.044(0.888) |
| Heatmap corr. | ASD | 0.671(*) | 0.607(*) | 0.528(0.007) | 0.460(0.034) | 0.379(0.125) | 0.333(0.208) | 0.261(0.424) |
|  | TD | 0.752(*) | 0.722(*) | 0.651(*) | 0.573(0.002) | 0.483(0.021) | 0.427(0.061) | 0.338(0.230) |
|  | TD-ASD | 0.081(0.500) | 0.114(0.460) | 0.124(0.557) | 0.113(0.678) | 0.104(0.753) | 0.094(0.814) | 0.077(0.888) |

**Table 3.**
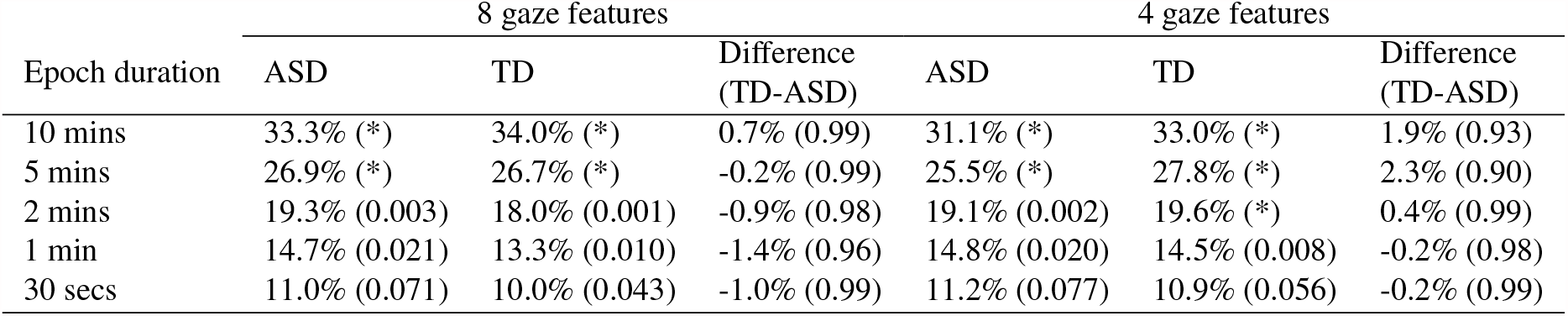
The change in fingerprinting analysis identification accuracy as a function of resampling epoch duration. The analysis procedure was performed either using eight gaze features (the percentage of time spent looking at screen, faces, non-social content, non-head body parts, hands, eyes, mouth, and heatmap correlations with TD-reference heatmaps) or four features (percentage of on-screen, face- and eye-looking time, heatmap correlations). Accuracy values were computed using two randomly sampled epochs of videos (duration given in columns) and then averaged across 10,000 resampling iterations. Values in parentheses show significance level (bootstrap test, FDR corrected for multiple comparisons within each epoch duration). Asterisk denotes *p <* 0.001.

## Notes

### Competing Interest Statement

The authors have declared no competing interest.

